# Experimental and mathematical approaches to quantify recirculation kinetics of lymphocytes

**DOI:** 10.1101/268326

**Authors:** Vitaly V. Ganusov, Michio Tomura

## Abstract

One of the properties of the immune system that makes it different from nervous and endocrine systems of mammals is the ability of immune cells to migrate between different tissues. Lymphocytes such as T and B cells have the ability to migrate from the blood to secondary lymphoid tissues such as spleen, lymph nodes, and Peyer’s patches, and then migrate back to the blood, i.e., they can recirculate. Recirculation of lymphocytes has been a subject of intensive investigation decades ago with wealth of data on the kinetics of lymphocyte recirculation available. However, these data have not been widely used to estimate the kinetics of recirculation of different lymphocyte subsets in naive and immunized animals. In this paper we review pioneering studies addressing the question of lymphocyte recirculation, overview quantitative approaches that have been used to estimate the kinetics of lymphocyte recirculation, and provide currently published estimates of the residence times of resting lymphocytes in secondary lymphoid tissues of mammals.

## Introduction

Adaptive immune system of mammals includes two major subsets of lymphocytes, B and T lymphocytes [1, 2]. One of the major functions of the adaptive immune system is to protect its host from invading microorganisms such as viruses and bacteria. Because microbes can enter the host from multiple areas such as via skin, lung or gut mucosa, there is a need to have lymphocyte being present near these tissues. In addition, because every lymphocyte is in general specific to only one microbial determinant (epitope) and there are many microbial determinants, only a small percent of lymphocytes would be able to recognize any specific microbe. For example, recent estimates suggest that only 1 in 10^5^ – 10^6^ of T lymphocytes would recognize a given epitope; that is in a mouse that has ∼2 × 10^8^ lymphocytes only about 200-2000 T cells would be specific to a given epitope [3]. While pathogens in general have multiple epitopes, only few of those can be recognized by lymphocytes. For example, in B6 mice lymphocytic choriomeningitis virus (LCMV) has about 30 epitopes that are recognized by CD8 T cells at all and only a few recognized strongly [4]. It is probably impossible to put a few thousands of T cells in all potential places of entry of a pathogen. Instead, lymphocytes have the ability to recirculate between different tissues in the body, thus increasing chances of encountering antigen they are specific for. This ability to migrate from the blood into several specific tissues and then back to the blood (i.e., to recirculate) makes adaptive immune system different from several other major systems of mammals such as nervous and endocrine systems. Because lymphocyte recirculation is a fundamental property of the mammalian immune system, our knowledge of how immune system works would be greatly incomplete if do not have solid understanding of the kinetics of lymphocytes recirculation, i.e., how quickly lymphocytes migrate to peripheral tissues, how long they spend in the tissues and return back to circulation. Understanding lymphocyte recirculation kinetics may be not just an academic exercise as blocking lymphocyte migration by anti-VLA4 antibodies – VLA4 is an integrin regulating lymphocyte entry into several tissues – has been shown to be effective in reducing symptoms of multiple sclerosis [5]. However, such treatment has serious side effects suggesting that deeper understanding how lymphocyte migration is regulated is needed [6, 7].

Ability of lymphocytes to recirculate between blood and tissues depends strongly on the type of lymphocyte, type of the tissue, and conditions of the host [8–18]. Specifically, naive T cells — cells that have not yet encountered their cognate antigen — are able to recirculate between blood and secondary lymphoid organs such as lymph nodes, spleen, and Peyer’s patches [9, 14, 19]. Activated T cells have the ability to migrate to nonlymphoid tissues [14, 20]; however, whether activated T cells in nonlymphoid tissues can migrate back to the blood remains poorly understood [21]. Inflammation may also change the pattern of lymphocyte migration; for example, intravenous (i.v.) infection may lead to trapping of recirculating lymphocytes in the spleen [22]. In this review we will focus on aspects of recirculation of resting (naive and memory) T lymphocytes with the major focus on migration of these cells via secondary lymphoid tissues, and how mathematical modeling has helped so far to quantify kinetics of this recirculation. Our main focus on recirculation of resting T cells is due to lack of good quantitative data and mathematical models on recirculation of activated T and B cells. However, we will provide a novel analysis of older data on recirculation kinetics of activated T cells in mice.

Because this review is about lymphocyte recirculation it is important to outline some basic anatomical features of the mammalian immune system. Since mice are the smallest mammalian animal model used to study lymphocyte recirculation, we focus our description specifically on murine secondary lymphoid organs. The major secondary lymphoid organs of mice are lymph nodes (LNs), spleen, and Peyer’s patches (PPs). Fluids that leak out of blood vessels are collected by the lymphatic vessels which bring this interstitial fluid via afferent lymphatics to tissue-draining lymph nodes. Each lymph node drains fluids from specific tissues and fluids (lymph) exit lymph nodes via efferent lymphatics often into another lymph node [23, 24]. Lymph from the final lymph nodes is collected into two big vessels, left and right lymphatic ducts, which are connected to the blood and which return collected interstitial fluids and cells back to circulation [2]. In mice and humans, the right lymphatic duct collects lymph from the upper right part of the body (about 1/4 of all interstitial fluid) and left lymphatic duct (also called thoracic duct) collects lymph from the rest (about 3/4) of the body. Lymph nodes can be roughly divided into several groups such as skin-draining lymph nodes, lung-draining lymph nodes, and gut-draining lymph nodes. A typical laboratory mice strain has about 30 lymph nodes [23] while humans have hundreds (perhaps over a thousand) lymph nodes [24–26]. Peyer’s patches are lymph node-like structures found in the gut. Peyer’s patches do not have afferent lymphatics and efferent lymph from Peyer’s patches flows into mesenteric (gut-draining) lymph nodes. Finally, spleen is probably the largest single secondary lymphoid organ in mice and humans [25, 27–31]. Spleen is not connected directly to the lymphatic system and lymphocytes enter the spleen from the blood and exit the spleen into the blood. In contrast, lymphocytes may enter lymph nodes or Peyer’s patches from the blood via high endothelial venules, and lymphocyte may also enter lymph nodes by migrating from the blood to peripheral tissues such as skin or gut, and then enter lymph nodes with afferent lymph. Thus, lymphocytes have several different pathways for recirculation in the body.

## Mathematical modeling of lymphocyte recirculation

By definition recirculation of lymphocytes is a dynamic process and therefore mathematical modeling is likely to be a useful tool for understanding of lymphocyte recirculation. Mathematical modeling is required to accurately quantify the kinetics of lymphocyte recirculation and to estimate the rates of lymphocyte migration from the blood to tissues and lymphocyte residence times in tissues. There have been many uses of mathematical models to understand cell migration. For example, mathematical models have been used to gain insights of how lymphocytes move in lymphoid and nonlymphoid tissues and how tissue composition impacts movement patterns of T cells [32, 33]. Here our main focus will be on experimental studies providing quantitative data on lymphocyte migration between tissues, and on mathematical modeling attempts to quantify lymphocyte recirculation kinetics (Table 1).

**Table 1:** Summary of published estimates of residency times of resting lymphocytes in secondary lymphoid tissues such as spleen, lymph nodes, and Peyer’s patches. Here ELL are efferent lymph lymphocytes (lymphocytes isolated by cannulation of individual lymph nodes, e.g., in sheep), TDL are thoracic duct lymphocytes (lymphocytes isolated by thoracic duct cannulation). In most other cases lymphocytes were isolated from lymph nodes and/spleen. Listed values for the lymphocyte residency time are as has been reported by authors and in some cases, half-life time of lymphocytes in the tissue (*T*_1/2_) was converted to the residency time using formula *T* = ln 2 × *T*_1/2_. In some studies, estimates for the residence time of lymphocytes in lymph nodes were dependent on the lymph node type (e.g., [73]), therefore, the presented estimates are for the pooled data. For time-dependent residency times the initial value was used (e.g., [71]).

| Lymphocyte type | Tissue/organ | Animal species | Residency time $T$, $h$ | Reference |
| --- | --- | --- | --- | --- |
| TDL | Spleen | Rats | 4.5 | [46] |
| TDL | Spleen | Rats | 6.0 | [66] |
| TDL | Spleen | Rats | 1.4 | [48] |
| TDL | Spleen | Rats | 2.4 | [49] |
| Blood lymphocytes | Spleen | Pigs | 3.0 | [28] |
| Naive T cells | Lymph nodes | Mice | 4.5 | [69] |
| naive CD8 T cells | Lymph nodes | Mice | 6.0 | [71] |
| memory CD8 T cells | Lymph nodes | Mice | 6.0 | [71] |
| naive CD8 T cells | Lymph nodes | Mice | 16.0 | [49] |
| memory CD8 T cells | Lymph nodes | Mice | 9.0 | [49] |
| naive CD4 T cells | Lymph nodes | Mice | 12.0 | [73] |
| naive CD8 T cells | Lymph nodes | Mice | 21.0 | [73] |
| CD4 T cells | Lymph nodes | Mice | 18.0 | this work |
| CD8 T cells | Lymph nodes | Mice | 18.0 | this work |
| B cells | Lymph nodes | Mice | 28.0 | this work |
| TDL | Lymph nodes | Rats | 20.0 | [66] |
| TDL | Lymph nodes | Rats | 10.0 | [49] |
| ELL | Lymph nodes | Sheep | 31.0 | [74] |
| TDL | Peyer's patches | Rats | 10.0 | [49] |

Kinetic aspects of lymphocyte recirculation has been studied since 1950^th^ using multiple mammalian species such as mice, rats, sheep, and pigs (see more below). Prior to pioneering experiments by Gowans the role of small lymphocytes found in the blood was unknown. Collecting lymphocytes from the thoracic duct lymph of rats over several days led to a decline in the number of lymphocytes found in the lymph. However, return of cells collected by thoracic duct cannulation back into circulation prevented loss of lymphocytes from the lymph [34]. These key experiments thus established that lymphocytes are able to recirculate between blood and thoracic duct lymph. By labeling lymphocytes collected from the blood or lymph (e.g., by thoracic duct (rats) or individual lymph node (sheep, pigs) cannulation) with radioactive labels further experiments demonstrated that indeed lymphocytes migrate from the blood to the efferent lymphatics of lymph nodes [8, 35–40].

### Spleen

Spleen is a large secondary lymphoid organ and it was previously estimated that about 20% of all lymphocytes in a human are found in the spleen [25, 31]. In mice, around 50% of all lymphocytes from secondary lymphoid tissues are found in the spleen [41, 42]. While it has been estimated that many lymphocytes travel via the spleen of mice, rats, or pigs [28, 30, 43] the exact time lymphocytes spend in the spleen has not been accurately quantified. Previous studies documented accumulation of radioactively labeled lymphocytes in the spleen after i.v. infusion of such cells [43–45] or dynamics of labeled cells in the blood in normal or splenectomized pigs [29]. However, data from these studies have not been analyzed using mathematical models, and thus, these previous data did not lead to estimates of lymphocyte residence (or dwell) times in the spleen.

An interesting approach was taken by Ford who designed an apparatus allowing maintenance of viable rat spleens for an extended period of time (up to 10 days, [46]). By creating an artificial circulation system connecting the spleen’s blood vessels, the author could monitor concentration of lymphocytes that exit the spleen during the perfusion of labeled thoracic duct lymphocytes that had been injected into circulation. Similar experiments with isolated pig spleens was done later by another group [28]. Verbal analysis of the data on migration thoracic duct lymphocytes via isolated perfused spleen led to estimate of lymphocyte residence time in the rat spleen of 4-5 hours [46, 47] and in the pig spleen of 2-4 hours [28]. A relatively complex mathematical model was proposed and fitted to the data of Ford [46]. This mathematical model-based analysis predicted that only about 10-25% of lymphocytes migrating via spleen pass via the marginal zone of the spleen (i.e., enter the spleen parenchyma). Cells entering the marginal zone of the spleen had a residency time of 50 minutes in the tissue. About 10% of cells existing the marginal zone migrated to the white pulp where the residency time was 4.6 hours. In contrast, remaining 90% of lymphocytes exited marginal zone into the red pulp with the average residency time of 5 minutes in that subcompartment of the spleen [48]. While Hammond [48] did not calculate the average time lymphocytes spend in the spleen, given the estimates provided, the average residency time of TDLs in the spleen is 50 + 0.1 × 4.6 × 60 + 0.9 × 5 = 82 min or 1.4 hours. This is a significantly shorter residency time than concluded by Ford [46] by visual analysis of the data.

Ganusov and Auerbach [49] used another set of data on migration kinetics of thoracic duct lymphocytes (TDLs) in rats. In these experiments (see also below), lymphocytes collected via thoracic duct cannulation were transferred into syngenic rats and the accumulation and loss of transferred cells (labeled with a radioactive label) were measured in multiple tissues of rats [50]. By fitting a mathematical model to these experimental data, Ganusov and Auerbach [49] estimated recirculation kinetics of TDLs including residence of these cells in major secondary lymphoid tissues of rats. The model fits predicted that average residence time of TDLs in the spleen is 2.4 hours which is also shorter than a previous estimate given in Ford [46] but longer than the estimate obtained from parameters of Hammond [48].

### Lymph nodes

Migration of lymphocytes from blood to lymph nodes and then back to the blood has been the main focus of many studies on lymphocyte recirculation. In part, this stems from the fact that in small animals such as rats cells that had migrated via lymph nodes can be easily collected via the thoracic duct cannulation, and, thus, the tempo of lymphocyte movement from the blood to thoracic duct can be easily recorded. In larger animals an efferent lymphatic vessel exiting a given lymph is wide enough to allow cannulation and collection of cells exiting that specific lymph node.

As described above, following the classical studies by Gowans illustrating ability of lymphocytes to recirculate between blood and thoracic duct lymph, there have been multiple studies investigating details of lymphocyte migration from blood to efferent lymph (reviewed in [30, 51–53]). Transit times via a particular lymph node (e.g., inguinal or cervical) have been inferred by transferring labeled lymphocytes into the animal and measuring rate of exit of labeled lymphocytes via a cannulated node in large animals such as sheep and pigs (e.g., [8, 9, 16, 54–57, Figure 1A]). Alternatively, kinetics of lymphocyte migration from the blood to the thoracic duct (i.e., via the whole lymphatic system) have been measured in smaller animals such as mice and rats (e.g., [36, 50, 58–65, Figure 1B]).

**Figure 1:**
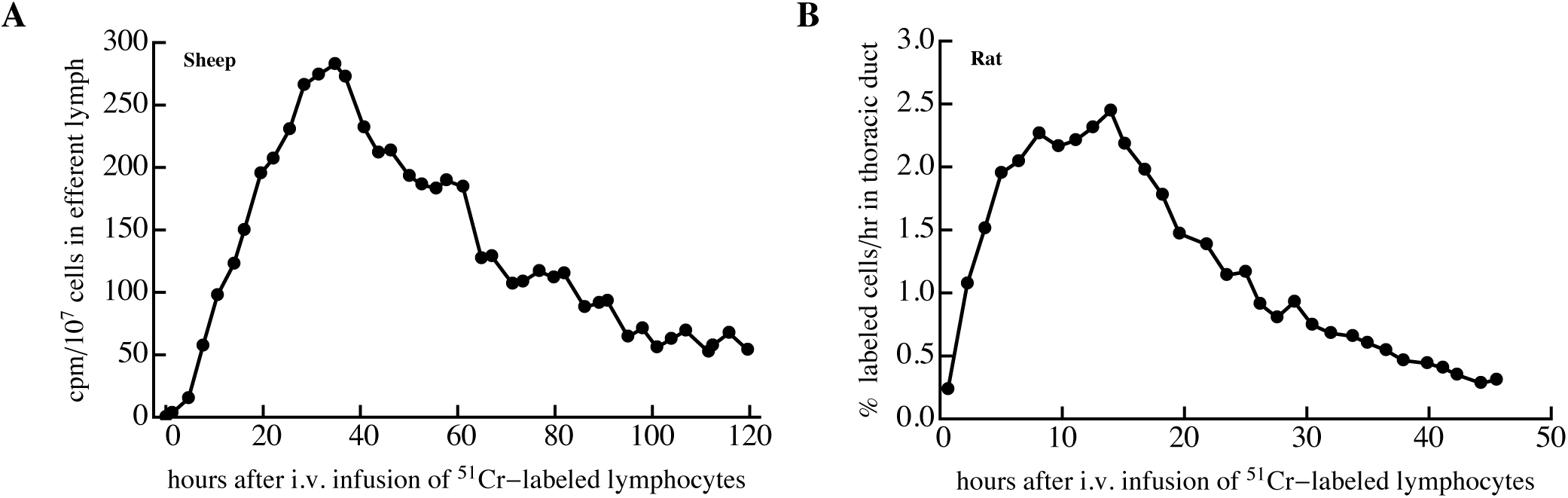
Typical examples of experimental data on lymphocyte migration from blood to lymph. In such studies, lymphocytes were isolated from a tissue (e.g., efferent lymph, lymph nodes, blood, spleen) and labeled with a label. In older studies labels were radioactive while newer studies involved fluorescent labels. Then cells were transferred intravenously into the same (e.g., sheep, panel A) or syngenic (e.g., rat, panel B) host. The accumulation of labeled cells in the efferent lymph of a given lymph node (panel A) or in the thoracic duct (panel B) were then followed. Different measures of the number of labeled cells have been used such as the number of labeled cells per fixed number of total isolated cells (e.g., estimated by measuring counts-per-minute (cpm) from the sample, panel A) or as the number or percent of injected cells (measured by total radioactivity) per unit of time (panel B). Data were digitized from previous publications (panel A: Frost et al. [54], panel B: Smith and Ford [50]).

While the data on the migration kinetics of lymphocytes via individual lymph nodes or from the blood to thoracic duct have been collected (e.g., Figure 1) very few studies attempted to estimate the lymphocyte residence (or dwell) time in lymph nodes from these cannulation experiments. For example, by simply looking at the data it is unclear which characteristic of the distribution observed in Figure 1 represents the average residence time. Mode, median, and average could all potentially be good estimates of the average residency time, and intuitively the average of the distribution has been treated in experimental studies as an estimate for residence time of lymphocytes in lymph nodes, e.g., about 48 hours in sheep or 24 hours in rats [30, 58]. However, to accurately estimate the average residence time one need to use mathematical modeling that takes physiology of the recirculatory and lymphatic system of mammals into account.

As far as we know the first mathematical modeling-based attempt to quantify lymphocyte migration via lymph nodes was in a series of papers by Stekel et al. [66–68]. The main idea of the mathematical model considered in these papers was the ability of lymphocytes to attach to and deattach from the lymphoid tissues while in lymph nodes or the spleen [66]. Deattached cells move through the tissue and this “movement” was described by a transport equation. Attached cells, however, would not move, and thus the process of “attachment – detachment” generated a skewed distribution and matched data on thoracic duct cannulation in rats [66, 67]. This work suggested 20h residency time of lymphocytes in the lymph nodes and 6 h residency time in the spleen of rats [66]. The model was further used to explain different kinetics of lymphocyte migration in irradiated or thymectomized rats [67, 68]. It is now well understood that the basic assumption of the model that cell attachment to structures in lymph nodes prevent lymphocyte exit is incorrect; rather, lymphocyte use structures including fibroblastic reticular cells to move around and to exit lymph nodes although precise mechanisms regulating lymphocyte exit from lymph nodes remain to be fully defined [69]. By using intravital imaging of T lymphocytes moving in murine lymph nodes and by mathematical modeling of T cell movement in the nodes Grigorova et al. [69] estimated the half-life time of T cells in the lymph nodes of mice to be 4-5 hours. Another study analyzed importance of directional movement of T cells in lymph nodes using digital reconstruction of a rat lymph node but the actual residency times of lymphocytes in the nodes was not estimated [70].

An important study quantified the residence time of antigen-specific naive and memory CD8 T cells using a novel “transfer-and-block” technique [71]. In this approach, naive or memory CD8 T cells, specific to the GP33 epitope of LCMV were transferred into congenic hosts and 24 hours after cell transfer further entry of lymphocytes into LNs was blocked by using anti-CD62L antibodies. CD62L is expressed on T cells and is generally required for cell entry into LNs [72]. The declining number of LCMV-specific T cells remaining in the LNs at different times after the blockade was used to infer the lymphocyte residence time. Interestingly, the authors found a non-monotonic rate of loss of T cells from LNs; naive and memory CD8 T cell populations had initial residency times of 5-6 h in LNs which increased significantly at later times for both naive and memory T cell populations to 15-16 hours [71]. Re-analysis of these data in another study revealed that the data could be accurately explained by a model in which residency time of lymphocytes is density-dependent and declines with time since blockade (if blockade was 100% efficient) [49]. In this reanalysis, naive and memory CD8 T cells had different residency times (16 and 9 hours for naive and memory T cells, respectively) [49]. Thus, the use of the same data but different assumptions on lymphocyte migration may result in different estimates of lymphocyte recirculation kinetics.

Mandl et al. [73] extended the study of Harp et al. [71] by transferring polyclonal naive CD4 and CD8 T cells into congenic mice and by blocking entry of new cells 2 hours after cell transfer by using a combination of antibodies to CD62L and VLA4. The authors found that a residence time of naive T cells was dependent on the type of the cell (CD4 vs. CD8 T cell). In contrast with previous work [71] the authors observed that after the blockade the percent of transferred cells declined exponentially over time. By fitting a line to log-transformed cell frequencies the authors estimated that naive CD4 and CD8 T cells spend on average 12 and 21 hours, respectively, in lymph nodes in mice [73].

It is interesting to note that we only know of one study that estimated residence time of lymphocytes from data generated by cannulating individual lymph nodes in sheep [74]. The authors proposed that lymphocyte migration within a lymph node can be described as a Markov process with *n* states and probability of jump from one state to another forward (*i* → *i* + 1) or backward (*i* → *i* – 1). Reaching the *n*^*th*^ state implied exit of a lymphocyte from the lymph node. The model was fitted to the cannulation data similar to that in Figure 1A and predicted the average residence time of lymphocytes in sheep lymph nodes of 31 hour [74]. Using a different mathematical model, which incorporated lymphocyte recirculation kinetics in the whole body, we have analyzed similar data on lymphocyte migration via individual lymph nodes in sheep (McDaniel and Ganusov (in preparation)). Depending on the dataset we found the average residence time of blood-derived lymphocytes in ovine lymph nodes to be 18-22 hours. This is another demonstration that the estimate of the lymphocyte residence time may not be robust to the choice of a model [75].

We recently performed mathematical modeling-assisted analysis of experimental data on recirculation of thoracic duct lymphocytes (TDLs) in rats [49]. In these experiments [50], lymphocytes collected in rats via thoracic duct cannulation were transferred into a series of syngenic hosts and accumulation and loss of the transferred cells in multiple lymphoid tissues was followed over time. By fitting a series of mathematical models to experimental data we estimated the TDL residence time in multiple tissues including lymph nodes. Interestingly, in contrast with previous studies that found differences in T lymphocyte residence times in different lymph nodes (e.g., mesenteric vs. subcutaneous LNs) [71, 73] we found the average residence time in subcutaneous or mesenteric LNs or PPs to be 10 hours [49, see more below].

**Figure 2:**
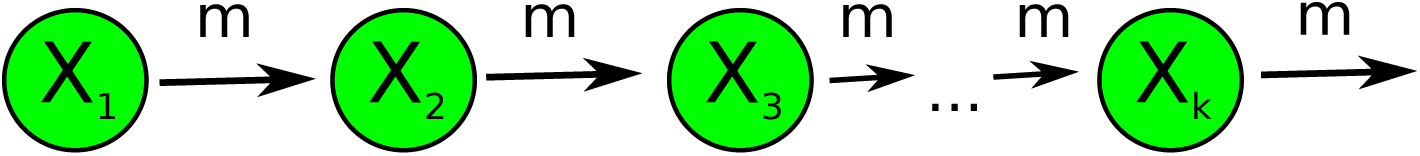
Schematic of the mathematical model describing loss of photoconverted cells from a LN of Kaede mice. We assume that cells entering a LN must undergo *k* exponentially distributed transitions before they can exit the node. In essence, this assumes that residency time of lymphocytes in a lymph node follows a gamma distribution. In the model *X*_*i*_ is the number of lymphocytes residing in the *i*^*th*^ compartment and *m* is the rate of transition of cells between subcompartments (see also eqns. (1)–(2)). Cells exiting the last subcompartment exit the lymph node into efferent lymph.

A novel experimental technique, the Kaede mice, allows to accurately track exit of lymphocytes from a given tissue such skin or a lymph node [17, 76]. Kaede mice express a photoconvertable protein which upon exposure to violet light changes color from green to red [76]. This unique system allows to label cells in one location, for example, an inguinal lymph node (iLN) or skin, and track the movement of labeled cells to other tissues in the body [17, 77]. We developed a simple mathematical model to track the dynamics of photoconverted (red) lymphocytes in the iLNs of Kaede (or other photoconvertable mice, e.g., KikGR [78, 79]) mice. The model assumes that cells exit the LN and are not able to re-enter the same LN. Experiments have shown that labeling of cells in the iLN distribute between all LNs and the spleen with approximately of 2-3% of cells in LNs being red [76]. Therefore, until the percent of photoconverted cells in the iLN is above 8-10%, re-entry of such cells into the node can likely be neglected if we assume that photoconversion and surgery associated with it do not induce strong inflammation impacting cell migration. In experiments, the lack of inflammation was recorded by similar size of iLNs prior and after the photoconversion.

Because our previous study suggested that distribution of residence times of TDLs in LNs was best described by a gamma distribution (and not by exponential distribution) [49], for experiments with Kaede we describe exit of lymphocytes from a LN as cell “migration” via multiple (*k*) subcom-partments in the LN:

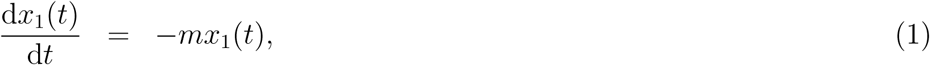

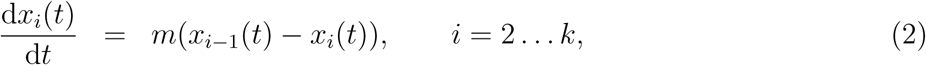

where *m* is the migration rate via a given subcompartment. The average residence time of lymphocytes in the lymph nodes is then given by *T* = *k*/*m*. Given that following photoconversion cells in all subcompartments at the steady state have equal densities, *x*_*i*_(0) = 1/*k*, the mathematical model (eqns. (1)–(2)) has a unique solution for the total number of cells in the LN 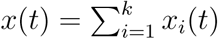:

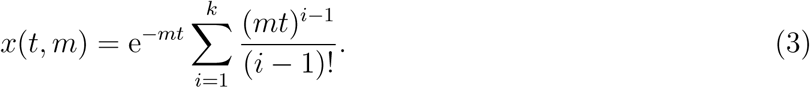

By fitting this solution (eqn. (3)) to experimental data from photoconversion experiments (Figure 3) we found that this simple model describes well the data for the loss of photoconverted CD8 T cells and B cells from the iLN (Figure 3B&C). However, the model was inadequate at describing the data for CD4 T cells as judged by the lack of fit test [80, results not shown]. Visually, this is likely because the loss of photoconverted CD4 T cells from the iLN is not exponential (compare Figure 3A & 3B). These results were independent of the number of subcompartments tested. While it is clear that CD4 and CD8 T cell populations most likely consist of subpopulations with perhaps different rates of exit from the iLNs, e.g., naive and memory T cells, why we were not able to detect such heterogeneity for CD8 T cells was unclear. It is possible that heterogeneity in CD4 T cells may come from a more diverse sets of cell types present in this group, for example, naive, memory, and regulatory T cells. Previously it was noted that naive and memory phenotype CD4 T cells have different exit kinetics from LNs [17]. In contrast, LCMV-specific naive and memory CD8 T cells appear to exit iLNs with similar kinetics [71].

**Figure 3:**
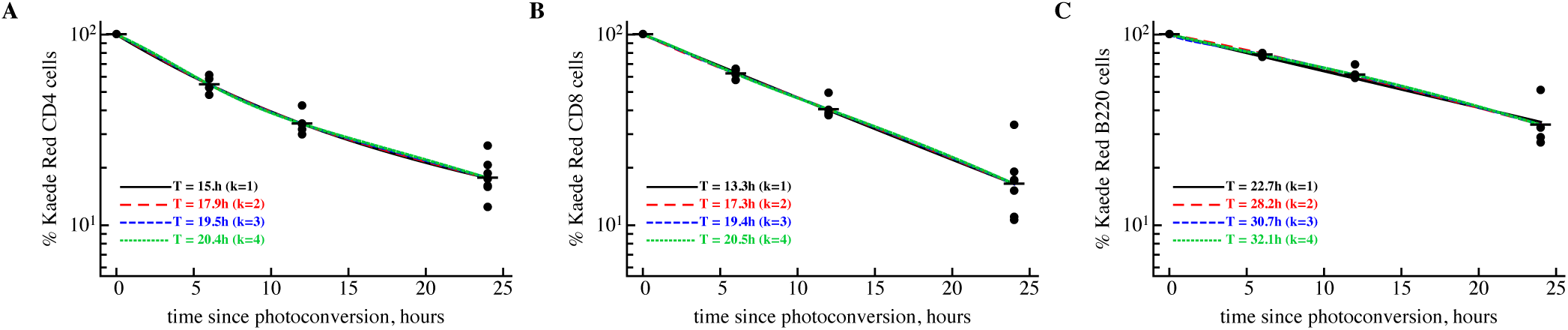
Estimated residence time of lymphocytes depends on the shape of residence time distribution. Lymphocytes in inguinal lymph nodes (iLNs) of Kaede mice were photoconverted [76] and the percent of CD4 T cells (panel A), CD8 T cells (panel B), or B cells (panel C) remaining in the iLNs at different times after phoconversion was recorded; the data from individual mice are shown by dots and horizontal lines denote average percent per time point. We fit a series of mathematical models assuming one (eqn. (3)) or two (eqn. (4)) subpopulations of cells with a different number of subcompartments (*k* = 1 … 4) to these data. Fits of the models with 2 subpopulations are shown by lines. To normalize residuals we used log_10_ transformation. The model with two cell subpopulations only improved the fit of the data for CD4 T cells (F-test, *p* < 0.005). Data for CD8 T and B cells were well described by a model with one, homogenous population (F-test, *p* > 0.05). Parameters of the model with 2 subpopulations and *k* = 2 compartments and their 95% confidence intervals (found by bootstrapping residuals with 1000 simulations) for different lymphocyte types are: CD4 T cells: *m*_1_ = 0.30 (0.30, 0.31)/h, *m*_2_ = 0.066 (0.065, 0.067)/h, *f* = 0.53 (0.52, 0.53), *T* = 17.9 (17.9, 18.0) h; for CD8 T cells: *m*_1_ = 0.46 (0.43, 0.48)/h, *m*_2_ = 0.10 (0.10, 0.10)/h, *f* = 0.16 (0.15, 0.15), *T* = 17.9 (17.9, 18.0) h; B cells: *m*_1_ = 0.071 (0.071, 0.071)/h, *m*_2_ = 0.07 (0.07, 0.07)/h, *f* = 0.37 (0.37, 0.37), *T* = 28.2 (28.2, 28.2) h.

To more accurately describe the kinetics of loss of photoconverted CD4 T cells from iLNs we extended the simple model (eqn. (3)) by allowing 2 subpopulations with relative frequencies *f* and 1 - *f* and with different exit kinetics determined by the rates *m*_1_ and *m*_2_, respectively. In this model, the total number of photoconverted (red) cells in the iLN is then given by

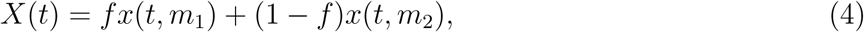

where *x*(*t, m*_*i*_) is given in eqn. (3). It is straightforward to extend this model to *n* subpopulations. The average residence time in this model is defined as *T* = *fk*/*m*_1_+(1 – *f*)*k*/*m*_2_ where *k* is the number of subcompartments in each of the subpopulations. This 2 subpopulation model can well describe experimental data on the loss of photoconverted CD4 T cells from iLNs (Figure 3A). Interestingly, for models that fit the data well (e.g., data for CD8 T cells or B cells) the estimated average residence time was not strongly dependent on the number of subpopulations assumed (1 or 2 subpopulations, results not shown). However, the estimate of the average residence time was strongly dependent on the number of subcompartments assumed, and, as the consequences, on the shape of the distribution of residence times. For example, the model fits predicted average residence time for CD8 T cells to be *T* = 13.3 h for *k* = 1 or *T* = 22.9 for *k* = 5 (eqn. (3)) with moderate reduction in the quality of the model fit to data at higher *k* as judged by AIC (results not shown). Therefore, it appears that the estimate of the average residence time from photoconversion data is not fully robust. Given our previous observation that best description of TDL recirculation kinetics via LNs in rats is given by a gamma distribution with shape parameter *k* = 2 our results suggest that average residence times in mouse iLNs are 18 h for CD4 and CD8 T cells, and 28 h for B cells.

### Whole body recirculation kinetics

An important limitation of many of the previously listed studies is that they considered migration of lymphocytes only via individual secondary lymphoid tissues such as spleen or individual lymph nodes. To study lymphocyte migration in the whole organism, Smith and Ford [50] adoptively transferred ^51^Cr-labeled TDLs and measured the percent of transferred lymphocytes in different organs of the recipient rats including the blood, lung, liver, spleen, skin-draining (subcutaneous) and gut-draining (mesenteric) lymph nodes. These data were initially analyzed with the use of a mathematical model [81] but TDL residence times in different tissues were not estimated. We developed a simple yet large mathematical model describing TDL dynamics, and by fitting the model to Smith and Ford [50] data, for the first time estimated the kinetics of TDL recirculation in the whole body [49, Figure 4].

**Figure 4:**
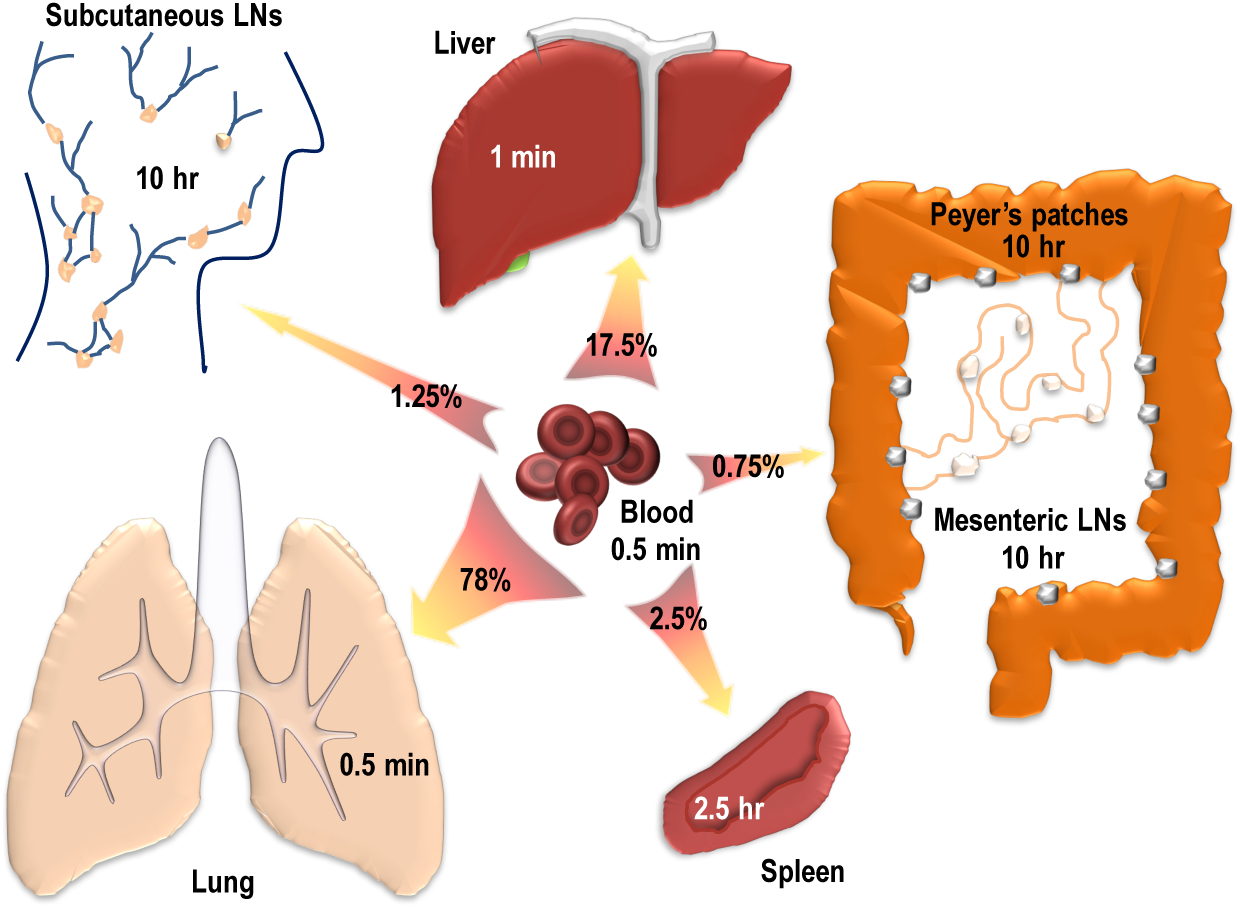
Estimated residence times of thoracic duct lymphocytes (TDLs) in rats [49]. We developed a mathematical model describing migration of lymphocytes in the whole organisms and by fitting the model to experimental data [50], estimated TDL residence times in major nonlymphoid and all major secondary lymphoid tissues of rats. Predicted residence times were short in the blood, lung and liver (*T* ≤ 1min) with 95% of lymphocytes recirculating between these tissues. Residence time in the spleen was 2.5 h with about 2.5% of lymphocytes entering the tissue. The remaining ∼ 2% of cells were entering lymph nodes and Peyer’s patches (PPs), with the average residence time of TDLs in these tissues was 10 hours [49].

The model fits predicted that TDLs spend very short time in the main blood vessels (about 30 sec) after which the vast majority of lymphocytes (about 95%) gets trapped in vasculature of the lung or the liver. This trapping is short-lived, however, and within 1 min trapped lymphocytes re-enter circulation (Figure 4). Only 5% of lymphocyte enter secondary lymphoid tissues per one passage of lymphocytes via circulatory system, with half of these entering the spleen, and half entering lymph nodes and Peyer’s patches (PPs). Lymphocytes reside for 10 h in LNs/PPs but only for 2.5 h in the spleen (Figure 4).

Because there is a good understanding of the kinetics of blood recirculation in rats, we would like to provide another interpretation of the kinetics at which TDLs pass via major tissues in rats. Previous studies found that the total blood volume in rats is proportional to the rat weight [82], and for 300 g rats, blood volume is *V*_*b*_ ≈ 20 mL. Heart volume is dependent on the animal size and age, and for 6-10 week old rats, *V*_*h*_ = 0.5 mL [83]. Given the high heart rate in rats (462/min) and stroke volume 0.3 mL per heart beat, the total cardiac output in rats is approximately *c* = 140 mL/min [84]. This in turn suggests the rate of blood recirculation in rats of *m*_0_ = *c*/*V*_*b*_ = 7/min or residency time of 9 seconds (i.e., on average in 9 seconds all blood passes through the heart). Given previously estimated rates at while TDLs enter various tissues [49], we predict that per one circulation of whole rat blood 27% and 6% of TDLs are attaching to the lung and liver vasculature, respectively. This suggests that the majority of TDLs pass via lung and liver vasculatures without attachment! Importantly, only 0.8% of TDLs in the blood will migrate to the spleen and 0.6% will migrate to LNs and PPs in one blood recirculation cycle lasting 9 seconds suggesting that the process of entering secondary lymphoid tissues by recirculating TDLs is not very efficient.

Differential probability of lymphocyte migration via lung and liver vs. secondary lymphoid organs is interesting but perhaps not unexpected given that these two organs are large and are expected to collect large volumes of blood. Amount of blood going to any specific tissue (cardiac output) has been measured in multiple species, for example, by measuring accumulation of labeled small microspheres after injection into the blood [40, 56, 84]. While we did not find a single study in rats measuring cardiac output to the same tissues as in our analysis (Figure 4), we found the cardiac output does not accurately predict the hierarchy of lymphocyte entry into lung, liver, and spleen. In rats, 0.7%, 3.3%, and 0.6% of cardiac output goes to lung, liver, and spleen, respectively [84], which is in contrast to 78%, 17%, and 2.5% of lymphocyte entry probability for these tissues [49]. Thus, migration and retention of lymphocytes in the whole body, while likely is dependent on the blood flow, is not strictly determined by the amount of blood going to any specific tissue.

### Recirculation of activated lymphocytes in mice

The vast majority of previous studies focused on quantifying migration of naive and memory lymphocytes via secondary lymphoid organs. While such lymphocytes are likely to represent the majority of cells in an organism in the absence of infection, infections will result in activation of lymphocytes. Yet, pattern and kinetics of migration of activated lymphocytes remain poorly defined. Cancer immunotherapy, involving *in vitro* expansion of populations of cancer-specific CD8 T cells and transfer of these cells into patients, is one of the novel ways to treat patients [85–87]. Therefore, deeper understanding of migration kinetics of activated T cells may help to improve the efficacy of T cell-based cancer therapies.

To determine the pattern and kinetics of recirculation of activated T lymphocytes we analyzed data from an old set of experiments [61]. In these experiments, Sprent [61] injected thymocytes (cells from the thymus) from CBA (H-2^*k*^) mice into irradiated CBA × C57Bl/6 (H-2^*k*^ × H-2^*b*^) F_1_ mice and isolated activated T cells via the thoracic duct cannulation [60]. Activated T cells were specific to the H-2^*b*^ antigen of the donor. Collected T cells were labeled with ^125^IUdR *in vitro* after 1 hour incubation and then injected intravenously into a series of syngenic CBA mice [61]. ^125^IUdR is incorporated into newly synthesized DNA, and therefore, only lymphocytes that were actively dividing *in vitro* became labeled.

**Figure 5:**
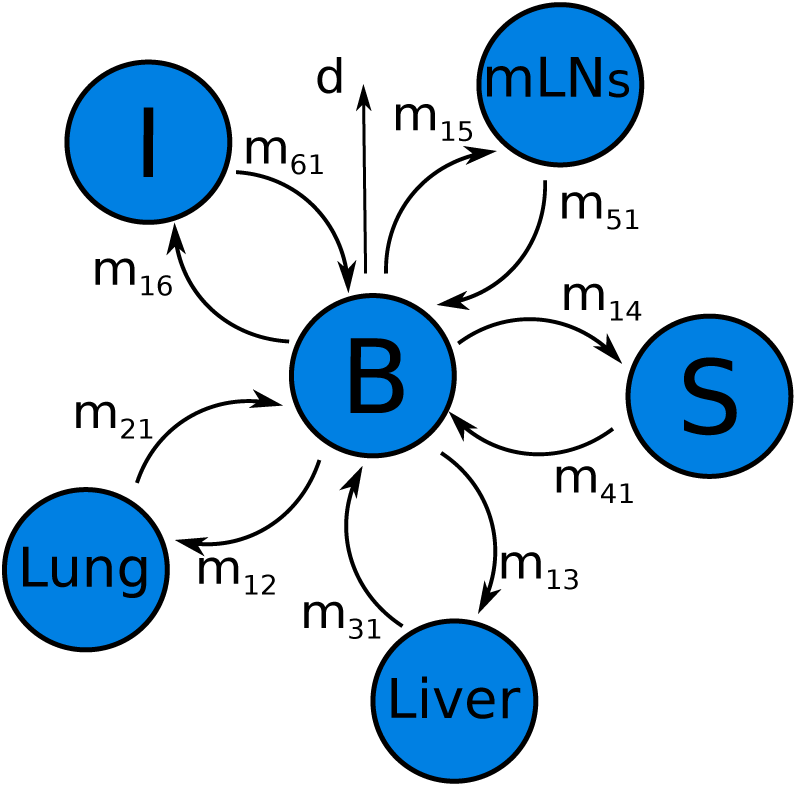
Schematic of assumed recirculation pathways of activated T lymphocytes [61]. In these experiments, lymphocytes were injected into the blood (*B*) and from the blood lymphocytes may enter lung (*i* = 2), liver (*i* = 3), spleen (*S, i* = 4), mesenteric LNs (mLNs, *i* = 5), intestine (*I, i* = 6) at rates *m*_1*i*_ with *i* = 2, … 6, respectively. Lymphocytes in these tissues may return to circulation at rates *m*_*i*1_ with *i* = 2, … 6. Lymphocytes may also leave the blood to other unsampled compartments and/or die at rate *d* (see eqns. (5)–(8)).

Following adoptive transfer recipient mice were sacrificed at different times after cell transfer and the percent of labeled lymphocytes was measured in several major organs of mice including blood, lung, liver, spleen, mesenteric lymph nodes (mLNs), thymus, kidney, and intestine [61, Figure 6]. Because very few cells migrated to thymus and kidney, we ignored these tissues in our following analysis; inclusion of these tissues did not influence significantly estimates of other parameters (results not shown).

**Figure 6:**
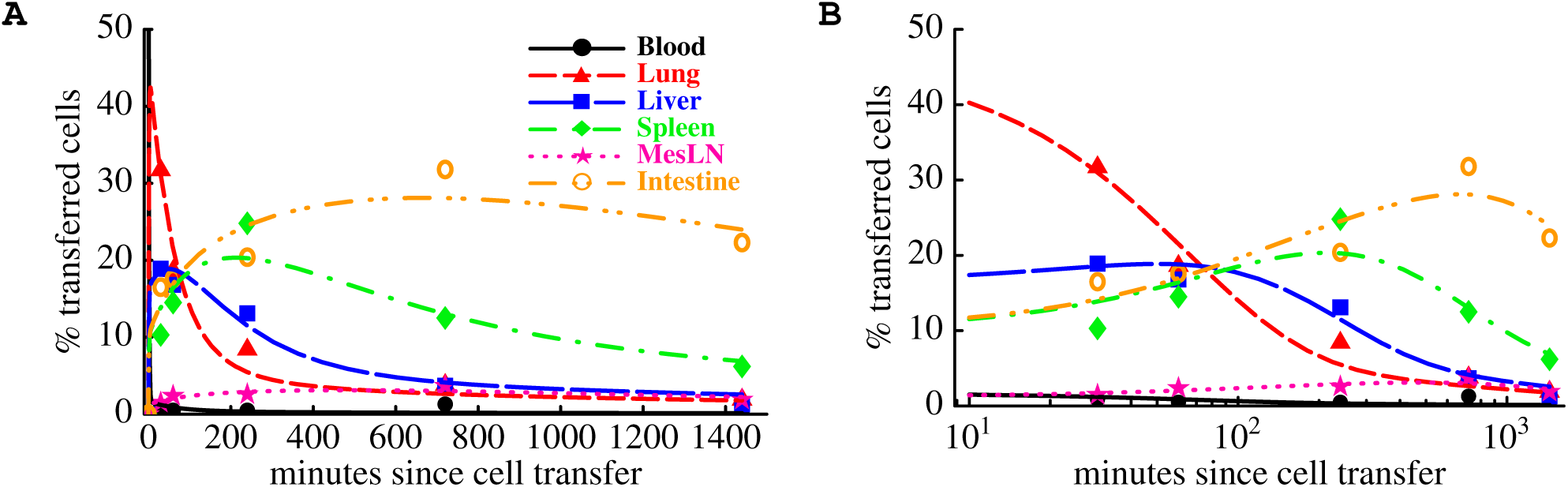
Experimental data and predictions of a mathematical model on the migration kinetics of *in vivo* activated T lymphocytes in mice. Thymocytes from CBA mice were transferred i.v. into CBA×C57BL/6 mice and activated TDLs were collected 4 to 5 days after cell transfer from the recipient mice. Replicating T cells were labeled with ^125^IUdR, adoptively transferred into CBA mice, and the distribution of transferred cells in different murine organs was followed over time (see Sprent [61] for more experimental detail). The percent of labeled lymphocytes recovered from different organs of the recipient mice are shown by markers. We fit the mathematical model of lymphocyte recirculation (eqns. (5)–(8)) to these experimental data; model fits are shown as lines. Plots are show on the linear (panel A) or a log-scale (panel B). Parameter estimates are given in Table 2.

To estimate the rates of activated T cell migration to major tissues of mice we adopted a mathematical model from our previous study [49]. In this model we assume that lymphocytes in the blood can migrate to multiple tissues such as lung, liver, spleen, and intestine, and following passage via the tissue, the cells would return back to the blood. The rate of lymphocyte entry into *i*^*th*^ tissue from the blood is denoted as *m*_1*i*_ and the rate of exit from the *i*^*th*^ tissue into the blood is then *m*_*i*1_ where *i* = 2, *…* 6. Following our previous work and some initial analyses in the model we assume that T cell migration via the lung and liver follows 1st order kinetics (i.e., is described by an exponential distribution), but residence in the spleen, mLNs and intestine is gamma-distributed. Gamma distribution of T cell residence in these tissues was modelled by assuming *k* subcompartments with migration rate *m*_*i*1_ between subcompartments. With these assumptions the mathematical model is given by a set of differential equations:

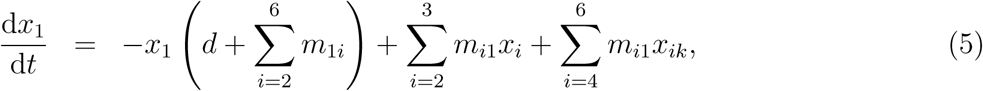

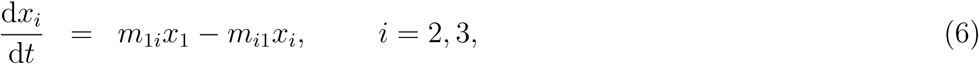

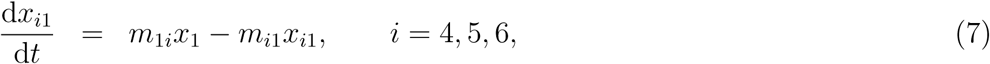

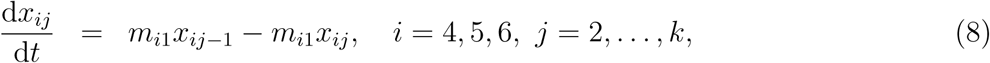

where *x*_*i*_ is the percent of labeled cells found in the blood (*i* = 1), lung (*i* = 2), liver (*i* = 3), and *x*_*ij*_ is the percent of labeled cells found in the *j*^*th*^ sub-compartment of spleen (*i* = 4), mesenteric LNs (*i* = 5), or intestine (*i* = 6), and *j* = 1 *… k, d* as the rate of removal of lymphocytes from circulation (due to death or migration to unsampled tissues such as other lymph nodes). Note that in this model we assume that cells migrating to the intestine return directly back to circulation without migrating via afferent lymph to mLNs. This assumption was justified by the lack of accumulation of labeled cells in the mLNs (Figure 6).

**Table 2:** Parameter estimates of the mathematical model that was fitted to the data on migration of *in vivo* activated TDLs in mice [61]. We list i) the rate of TDL entrance into a particular organ from the blood *m*_1*i*_ (second column), ii) the percent of cells leaving the blood into a particular organ 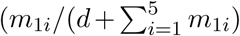, third column), iii) the rate of exit of TDLs from an organ to the blood *m*_*i*1_ (fourth column), and iv) the average residence time of TDLs in the organ (fifth column). The rate of migration of TDLs from the blood to all organs, 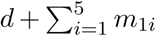, is 51.2 h^-1^ or the average residence time of cells in the blood is 1.2 min. The average residence time of cells in a particular organ is calculated as 1/*m*_*i*1_ (for blood, lung, and liver) and 2/*m*_*i*1_ for spleen, mLNs, and intestine. We assume *k* = 2 subcompartments in these latter organs since this allowed for the best description of the data based on AIC (results not shown). The rate of cell migration from the blood to other organs in the body *d* = 7.50 (3.68 - 10.02) h^-1^. No dying cells were assumed to migrate to the liver as including this process did not improve the quality of the model fit to data (not shown). In brackets we show 95% confidence intervals calculated by bootstrapping the residuals with 1000 simulations [88].

| Organ | Rate of entrance from blood $m_{1i}$, $\text{h}^{-1}$ | Percent cells going to organ from blood | Rate of exit from organ to blood $m_{i1}$, $\text{h}^{-1}$ | Residence time in organ, h |
| --- | --- | --- | --- | --- |
| Lung | 23.64 (13.3–44.3) | 46.1 (37.1–61.6) | 1.7 (1.0–3.6) | 0.6 (0.3–1.) |
| Liver | 8.61 (5.0–12.8) | 16.8 (12.1–20.6) | 0.4 (0.3–0.7) | 2.2 (1.3–3.3) |
| Spleen | 5.4 (2.9–7.7) | 10.5 (7.3–12.8) | 0.2 (0.2–0.4) | 8.0 (5.7–11.) |
| MesLN | 0.7 (0.3–1.6) | 1.3 (0.6–2.9) | 0.1 (0.0–0.8) | 16.8 (2.5–> 50) |
| Intestine | 5.5 (2.9–7.3) | 10.7 (7.2–12.8) | 0.1 (0.1–0.1) | 22.0 (15.4–31.8) |

The model was fit to experimental data using least squares. A number of interesting observations emerged. First, the model predicted a very short average residence time of activated lymphocytes in the blood, *T* ∼ 1.2 min (Table 2). Nearly 65% of activated lymphocytes in the blood migrated to the lung and liver where they spent on average 35 min and 2.2 hours, respectively. These residence times are substantially higher that those for resting TDLs [49, Figure 4]. It is interesting to note that trapping of activated lymphocytes in the liver after the i.v. injection has been observed previously in other experiments [22]. The spleen and intestine took another 20% of activated lymphocytes in the blood, and the average residence times were 8 and 22 hours in the spleen and intestine, respectively (Table 2). In these experiments very few cells migrated to the mesenteric lymph nodes and we could not reliably estimate the residence times of these *in vivo* activated lymphocytes in the mLNs (e.g., note large confidence intervals in Table 2 for this tissue compartment). We also found that about 15% of transferred cells in the blood per circulation cycle migrated to tissues/organs which have not been sampled in these experiments, e.g., skin or other lymph nodes, or these cells have been dying at a high rate. Indeed, it is expected that activated lymphocytes undergo programmed cell death following clearance of the antigen which has been mimicked by the adoptive transfer experiments. In 24 hours post-transfer, only 34% of injected radioactivity could be recovered from recipient mice [61, Figure 6 and results not shown]. Nevertheless, this analysis highlights a similar hierarchy of migration of naive and activated lymphocytes *in vivo* with the majority of cells entering lung and liver vasculature but residing there for a relatively short period of time in these organs as compared to other tissues.

## Recirculating and non-recirculating lymphocytes

Data and analyses presented so far may create an impression that all (or nearly all) lymphocytes of the immune system recirculate. This is not likely to be the case in general. Multiple factors are likely to influence ability of lymphocyte to recirculate following adoptive transfer [50, 89]. For example, it was noted that handling of lymphocyte *in vitro* at low temperatures dramatically impedes lymphocyte recirculation kinetics; passaging of lymphocytes via an intermediate host before final transfer into definite hosts restores the ability of lymphocytes to recalculate [89]. Accurate counting of lymphocytes entering isolated perfused spleens allowed to conclude that a large fraction of spleen lymphocytes never exit the tissue during 7-10 days of perfusion [46]. More recently, experiments involving surgical joining of syngenic mice (parabiosis) allowed accurate tracking recirculation kinetics of memory CD8 T cells in mice [90]. Initial studies showed limited ability of memory T cells in tissues such as brain to recirculate between two parabiotic mice [90]. A more thorough follow up revealed an inability of LCMV-specific memory CD8 T cells residing in most nonlymphoid tissues to recirculate between parabiotic mice [91]. Interestingly, some of such tissue-resident memory T cells were also found in secondary lymphoid tissues such as lymph nodes and spleen [92]. Because many of such non-recirculating T cells reside in peripheral, non-lymphoid tissues, perhaps, it is not totally surprising that they are not able to recirculate. However, non-recirculating cells could even be found in the blood, e.g., crawling along sinusoids in the liver [93, 94]. It is important to realize, however, that many of studies documenting “non-recirculatory” nature of lymphocytes have been performed for a relatively short-time period and deeper, mathematical modeling-assisted analyses of the data from such parabiosis experiments are needed in order to accurately quantify the residency time of such tissue-resident lymphocytes.

## Summary

There have been multiple studies documenting lymphocyte migration kinetics via secondary lymphoid organs of mice, rats, pigs, and sheep. Many studies involved simple or complex mathematical models to estimate residence times of lymphocytes in different tissues and in different conditions. A quick comparison reveals that these estimates while being approximately similar still vary dramatically in absolute values (Table 1). It is unclear at present if such variability in estimates is simply due to differences in experimental methodologies involved, in mathematical modeling approaches, or both. Future studies should attempt to determine whether estimates of the residence times are robust to the choice of mathematical model. It is often expected (and was found in this analysis) that conclusions arising from models being fitted to data, for example, estimates of model parameters, can be model-dependent [75]. It it also possible that there may not be such a universal parameter such as lymphocyte (e.g., naive CD8 T cell) residency time in a LN or spleen. Residency time may depend on the environment lymphocyte is in (resting vs. inflammed LN), previous history of the lymphocyte, or other factors. Future studies will have to become more mechanistic and instead of simply measuring/estimating lymphocyte migration kinetics, should attempt to determine why some lymphocytes spend 10 hours in tissues while other lymphocytes only 5 hours. Fundamental understanding of lymphocyte recirculation kinetics (or absence of thereof for tissue-resident lymphocytes) should allow to improve therapies that involve lymphocytes such as cancer immunotherapies.

## Acknowledgments

We would like to thank multiple people contributing to our discussions on lymphocyte recirculation including Reinhard Pabst, Jurgen Westermann, Dave Masopust, Gudrun Debes, Rob De Boer, Johannes Textor, Judith Mandl. We thank Jeremy Auerbach for the cartoon of the recirculation kinetics of thoracic duct lymphocytes. This work was supported by the NIH grant to VVG (R01 GM118553).

## Abbreviations

LNs: lymph nodes
iLNs: inguinal lymph nodes
mLNs: mesenteric lymph nodes
TDLs: thoracic duct lymphocytes
EEL: efferent lymph lymphocytes
LCMV: lymphocytic choriomeningitis virus
PPs: Peyer’s patches

